# Radioprotective role of cyanobacterial phycobilisomes

**DOI:** 10.1101/281386

**Authors:** Konstantin E. Klementiev, Eugene G. Maksimov, Danil A. Gvozdev, Georgy V. Tsoraev, Fedor F. Protopopov, Irina V. Elanskaya, Sergey M. Abramov, Maksim Yu. Dyakov, Vyacheslav K. Ilyin, Nadezhda A. Nikolaeva, Yury B. Slonimskiy, Nikolai N. Sluchanko, Victor M. Lebedev, Andrew V. Spassky, Thomas Friedrich, Georgy V. Maksimov, Vladimir Z. Paschenko, Andrew B. Rubin

## Abstract

It is now generally accepted that cyanobacteria are responsible for production of oxygen, which led to the so-called “Great Oxygenation Event”. Appearance of dioxygen in Earth’s atmosphere resulted in formation of the ozone layer and the ionosphere, which caused significant reduction of ionizing radiation levels at the surface of our planet. This event not only increased biological diversity but also canceled the urgency of previously developed mechanisms of DNA protection, which allowed to survive and develop in harsh environmental conditions including exposure to cosmic rays. In order to test the hypothesis if one of the oldest organisms on Earth retained ancient protection mechanisms, we studied the effect of ionizing radiation (IoR, here: α-particles with a kinetic energy of about 30 MeV) and space flight during the mission of the Foton-M4 satellite on cells of *Synechocystis* sp. PCC6803. By analyzing spectral and functional characteristics of photosynthetic membranes we revealed numerous similarities between cells exposed to IoR and after the space mission. In both cases, we found that excitation energy transfer from phycobilisomes to photosystems was interrupted and the concentration of phycobiliproteins was significantly reduced. Although photosynthetic activity was severely suppressed, the effect was reversible and the cells were able to rapidly recover from stress under normal conditions. Moreover, *in vitro* experiments demonstrated that the effect of IoR on isolated phycobilisomes was completely different from such *in vivo*. These observations suggest that the actual existence and the uncoupling of phycobilisomes under irradiation stress could play specific role not only in photo-, but also in radioprotection, which was crucial for early stages of evolution and the development of Life on Earth.

## 1. Introduction

It is well known that solar and cosmic radiation is lethal outside the ionosphere of the Earth, which is composed by ionized nitric oxide and dioxygen. Below the ionosphere, another thin atmospheric stratum, which is called ozone layer, blocks a significant amount of UV-C and –B light. Thus, di- and trioxygen essentially separate the surface of our planet from Space, but it has not always been like that. During the first two billion years of Earth’s existence, its atmosphere contained no oxygen. The radiation dosage arriving on the Earth’s surface exceeded today’s level by approximately 15 times for UV and about 46.6 times for IoR^1^. The situation changed after the appearance of photosynthetic organisms, which were able to use water as a source of electrons and evolved dioxygen as a side product. One must admit that this strategy was very successful, since the amount of oxygen produced by microorganisms was sufficient to oxidize every mineral on the Earth’s surface and to create the atmosphere^2-4^. Although the Great Oxygenation Event (GOE) took more than 900 million years to complete, it was a dramatic and game-changing incident, which affected biological diversity on our planet.

All modern geological and biological evidences indicate that communities of cyanobacteria are responsible for the GOE^5^, and it is often assumed that modern and ancient cyanobacterial species are quite similar. However, it is hard to believe that under different selection factors, namely in the presence and in the absence of UV-C, -B and IoR, these primitive organisms remained completely the same over billions of years of evolution since then. Here, we would like to focus on the selection factors, which were crucial for cyanobacterial development before the GOE.

First of all, oxygenic photosynthesis requires effective bioenergetic conversion of light into the energy stored in separated charges and chemically stable equivalents. The first step of this complex process implies absorption of a photon by antenna pigments and transfer of the excitation energy to the reaction center of the photosystem. Thus, antennas which transfer excitation energy (EET) to the photosystems significantly increase the effective absorption cross-section of the reaction center. In cyanobacteria, the light-harvesting antennas consist of huge water-soluble pigment-protein complexes, called phycobilisomes (PBs)^6-12^, the absorption of which in the visible spectral region is accomplished by linear tetrapyrroles, which are covalently bound to specific cysteines of phycobiliprotein subunits of the PBs. Different species of cyanobacteria utilize spectrally different types of phycobiliproteins^13-15^ with an overall quite similar structure as building blocks, which eventually compose a mega-Dalton complex harboring thousands of chromophores^16^. PBs are so huge that they could be easily visualized in an electron microscope, and in some species of cyanobacteria they reach up to 60% of total cellular protein and up to 20% of dry mass of the cell^17^. Considering the efficiency of cyanobacterial photosynthetic apparatus and the robustness of their survival strategy, which was evolved before the GOE, it may seem surprising that algae and higher plants completely lack PBs, although it is generally accepted that their chloroplasts had been evolved from cyanobacteria^18^ after endosymbiosis. This fact indicates that utilization of this giant for light-harvesting only was not energetically favorable for algae and plants, since evolution in eukaryotic photosynthesis developed membrane-embedded light-harvesting complexes instead. This immediately raises the question what else PBs were so necessary for, on top of light-harvesting in cyanobacteria?

Prior to the GOE, all living organisms were under pressure of other selection factors, and, in particular, protection of the integrity of DNA was vital. Several strategies were developed, including replication, storage of multiple copies of the same plasmid, chromosome, and genes^19^, but from these, prevention of DNA damage was probably the most effective. A general way to prevent excitation/ionization of DNA is to hide it behind some shield, which will be able to dissipate the energy into heat, to scavenge radicals, or to sacrifice itself in any other way for the greater good. This strategy probably became even more relevant due to the appearance of oxygenic photosynthesis, which significantly increases the probability of Reactive Oxygen Species (ROS) formation. Several examples of protective proteins consisting of hundreds of aromatic amino acids are known among different microorganisms. A particular (e.g., enzymatic) role of such proteins is unknown, however, UV-protection and/or involvement in storage of certain amino acids is possible. One such example is the S-layer carotenoid-binding protein of *Deinococcus radiodurans* (gene DR_2577), an organism that is known for its extreme ability to resist up to 200 times higher doses of IoR compared to *Escherichia coli*^20^. Thus, we can assume that the development of huge protein complexes like PBs was beneficial for both – radiation protection and light-harvesting, however, the last one probably was not mandatory. This hypothesis is supported by the fact that several *Synechocystis* mutants completely lacking PBs^21^ are able to grow autotrophically, however much more susceptible to environmental stress.

It should be noted that utilization of PBs also required the development of specific mechanisms which regulate their energetical coupling with photosystems. One such mechanism is called “state transitions”, which involves redox-dependent redistribution of PBs between the two types of photosystems^22-25^. Which molecular cascade triggers this reaction and how mobility of PBs is regulated is not sufficiently studied. Another mechanism requires so-called carotenoid proteins which can induce quenching of the excess of excitation energy in PBs, thus preventing EET to reach the photosynthetic reaction center^26-30^. In some species like *Synechocystis*, PBs quenching is triggered by photoactivation of the Orange Carotenoid Protein (OCP)^31,32^, however some species have multiple copies of full-length OCP-like genes and its fragments (Helical Carotenoid Proteins - HCPs^33,34^ and C-terminal Domain Homologs - CTDHs^35^) which can bind carotenoids, but not all of them are able to induce PBs quenching. Recent studies demonstrated that carotenoid molecules, which are known for their excellent antioxidant properties, could be transferred from membranes into water-soluble HCPs via CTDH, with the state of the latter being regulated by redox conditions^35-37^. The presence of multiple completely different water-soluble carotenoid carriers, which are not involved in direct PBs quenching, raises the question about their present and past functional roles and their involvement into photo- and other protection mechanisms. It should be noted that phycobiliproteins also have antioxidant activity, and, as reported in some pilot studies, can even stimulate radioprotection in lymphocytes from nuclear power plant workers^38^.

In this work we present a study about one of the few biological species, which were able to survive during Foton-M4 satellite mission^39^ - cells of *Synechocystis* sp. PCC6803 cyanobacterium. In order to deeper understand the observed effects of space flight, we conducted a series of *in vivo* and *in vitro* model experiments using accelerated α-particles as a source of IoR, which allowed us to suggest the radioprotective role of phycobilisomes.

## 2. Materials and Methods

### 2.1 Cultivation of Synechocystis sp. PCC 6803 and the Foton-M4 satellite mission

Wild type (WT) and mutants of cyanobacteria *Synechocystis sp.* PCC6803 from the collection of Biological Faculty of the Lomonosov Moscow State University were grown on a modified liquid medium BG-11 at 30 ° C under a white light lamp delivering 40 μmol photons m^-2^ s^-1^ to the surface of the reactor.

The experiment “BIORADIATION-F” was conducted within the framework of a collaboration of Lomonosov Moscow State University and the Institute of Biological and Medical Problems of the Russian Academy of Sciences. The tasks of this experiment were to study the biologically significant characteristics of cosmic IoR and the effects of its impact on biological objects in open space and satellite environments.

Prior sending to Baikonur cosmodrome (Kazakhstan), liquid cell culture of *Synechocystis* sp. PCC 6803 was inoculated into 5-ml sterile tubes poured with agar (0.6%) medium containing glucose and thoroughly mixed. Control samples were stored in the laboratory at 21 °C in the dark for the entire time of flight of the satellite mission. At the cosmodrome, samples were placed inside the satellite, into the thermostatted chambers maintaining temperature at +21 °C. The Foton-M4 satellite was sent into space on July 19, 2014 and spent 45 days on the orbit at the average height of 575 km. (http://biosputnik.imbp.ru/eng/Foton4.html). Temperature inside of the landing module was in the range from 14 to 22 °C^40^. At the beginning of the Foton-M4 mission, there was a communication failure, which has been restored after eight days. This incident resulted in a reduction of orbital mission duration. After landing of the satellite, samples were delivered to Moscow and were studied by various optical methods. Part of the culture which returned from the space mission was placed into liquid medium (BG-11) and cultivated under normal conditions. A similar procedure was carried out with the control samples kept in the laboratory.

### 2.2 Samples for in vitro experiments

PBs and Phycocyanin (PC) were isolated from the wild-type *Synechocystis* sp. PCC 6803 according to procedures described in^41^.

OCP with the N-terminal His-tag was expressed in a carotenoid-producing *E. coli* strain and purified as described in^41,42^.

### 2.3 Exposure to ionizing radiation

In order to model the effect of IoR by high (relativistic) energy elements (HZE) of cosmic rays, we used a 120-cm cyclotron developed at the Lomonosov MSU Skobeltsyn Institute of Nuclear Physics, which allows to obtain accelerated helium nuclei (α-particles) with energies up to 30.4 MeV. A beam of α-particles from a cyclotron was extracted from the ion guide through 50 μm thick aluminum window into the air and directed thought a replaceable diaphragm onto a sample in a 1.0 mm cell covered with 20 μm thick mylar film windows. The beam was monitored by the magnitude of the charge on the diaphragm and the cuvette with absolute accuracy of absorbed dose determination estimated as 30%, relative accuracy - not worse than 10%. The energy losses of α-particles at the “window” of ion guide, air layer, and mylar film were 4.2 MeV, so that the α-particles energy on the inner surface of the film was about 26.2 MeV. The linear energy transfer (LET) of α-particles with such energy is about 25 keV/μm in water and grows as the particles in the suspension slow down by approximately an order of magnitude. The LET value of the particles at the entrance to the sample chamber is close to the value of the LET of the relativistic neon-magnesium nuclei of galactic cosmic rays and then increases in the range, reaching 230 keV/μm corresponding to heavier nuclei (Si), which allows simulating the effect of heavy nuclei Solar Cosmic Rays and Galactic Cosmic Rays^43,44^. Since the mean free path of an α-particle with an energy of 26.2 MeV in water is equal to 0.45 mm, irradiation of the cuvette was carried out from both sides of the cell. The value of the absorbed dose was averaged over the entire volume of the cuvette and was equal to 30, 60, or 120 kGy.

### 2.4 Optical methods for the assessment of spectral and functional characteristics of photosynthetic complexes in vivo and in vitro

Fluorescence measurements were performed by time- and wavelength-correlated single photon counting system SPC-150EM with hybrid detector HMP-100-40 (Becker&Hickl, Berlin, Germany) as described in^45^. Excitation was performed with a pulsed 405 nm laser diode (InTop, St. Petersburg, Russia) and BDL-473-SMC (Becker&Hickl, Berlin, Germany) laser, delivering picosecond excitation pulses, driven at a repetition rates up to 50 MHz. Temperature of the sample was stabilized by a Peltier-controlled cuvette holder Qpod 2e (Quantum Northwest, Liberty Lake, WA). Fluorescence decay kinetics were approximated by the sum of exponential functions. Calculations were performed using SPCImage (Becker&Hickl, Germany) software packages.

Measurements of steady-state fluorescence spectra and fluorescence excitation spectra were carried out on a modified Fluorolog 3 (Horiba Jobin Yvon) spectrophotometer.

Absorption spectra were measured using a Lambda-25 spectrometer (Perkin Elmer).

Fluorescence induction curves were recorded using a PSI-3500Fl fluorimeter (PSI).

Variable fluorescence (Fv / Fm) was calculated according to procedures described in^46,47^.

All calculations were performed using the Origin 9.1 (OriginLab Corporation, USA). Each experiment except Foton-M4 satellite mission was conducted at least 3 times.

### 2.5 ROS-induced bleaching of pigments in vitro

Photosensitizing activity of polycationic aluminum phthalocyanines (AlPC) was used to study the effect of ROS on PBs and OCP. In order to induce ROS production, a mixture of AlPCs was illuminated by a 200 mW red LED with the emission maximum at 635 nm (Thorlabs, USA), which overlaps with the main absorption band of AlPC. In order to prevent photoactivation of OCP by the red LED, the sample was illuminated through a 600 nm longpass filter (Thorlabs, USA). Upon illumination of the sample, absorption in the visible range was recorded using a MayaPro spectrometer (Ocean Optics, USA).

## 3. Results

### 3.1 Absorption measurements reveal that PBs are sensitive to ionizing radiation

The main absorption bands of *Synechocystis* cells in the visible/near-infrared (Vis-NIR) region of the spectrum are attributed to pigments of the photosynthetic apparatus, namely chlorophyll *a* (Chl, main peaks at 440 and 680 nm) of the photosystems 1 and 2, carotenoids (shoulder at ∼ 500 nm) and phycobiliproteins of PBs, which are dominated by phycocyanin (PC, 623 nm). *Figure 1A* shows characteristic changes in absorption spectra of *Synechocystis* sp. PCC6803 cells after exposure to IoR and space flight. Upon an increase of the absorbed dose, we observed a decrease of optical density in the regions of absorption of both Chl and PBs. This effect was characterized by saturation for the Chl component and almost linear for the PBs component of the spectrum (*Figure 1B*). *In vitro*, comparable concentrations of isolated PBs were 50% bleached already at 60 kGy, which may indicate that *in vivo* PBs are more stable, or that the effect of ionizing radiation is variable between different cell compartments. Probably, due to the layer structure of the thylakoids, each layer shields the layer underneath. Such protection density might not be reached *in vitro*. In all experiments with cells we observed a decrease of the PBs-to-Chl absorbance ratio (*Figure 1C*), which indicates that PBs are more sensitive to IoR stress than Chl containing compartments. It should be noted that storage of cells in the dark, even in the presence of glucose resulted in reduction of PBs concentration, which is typical for different types of starvation^48,49^. Reduction of PBs absorption was even more pronounced in the cells, which traveled into space via the Foton-M4 satellite, in total resulting in a decrease of the PBs/Chl ratio comparable to the sum of the effects of starvation and exposure to IoR. It should be considered as a lucky coincidence that the duration of the space mission was shortened by two weeks. Otherwise, the longer starvation period could have eliminated the differences between the sample and the reference stored in the laboratory.

**Figure 1.**
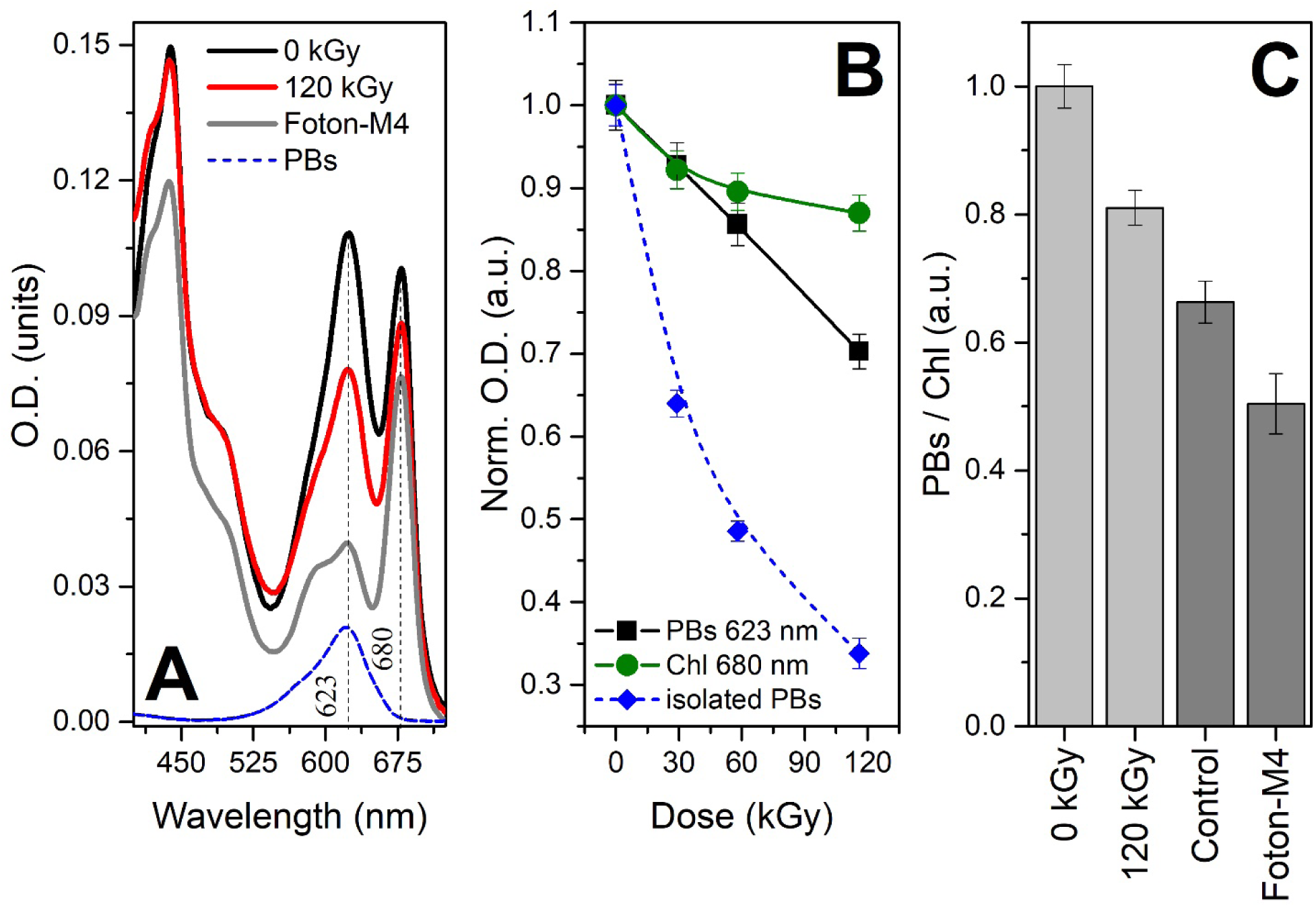
(**A**) - absorption spectra of *Synechocystis* sp. PCC6803 cells before and after exposure to 120 kGy of α particles (black and red lines, respectively) or the Foton-M4 space mission (gray, scaled). The blue dashed line shows the absorption of isolated PBs. (**B**) – dose dependency of optical density of *Synechocystis* sp. PCC6803 cells (PBs absorption, black; and Chl absorption, green) and isolated PBs (blue) from irradiation with α-particles. Absorption was normalized to O.D. values at 0 kGy. (**C**) - ratios of optical density of *Synechocystis* sp. PCC6803 cells at 620 nm (PBs) and 680 nm (Chl) after different treatments.

### 3.2 PBs and Chl fluorescence reveal that ionizing radiation affects energetic coupling of antenna and PS2

As we observed significant changes in concentrations of pigments in *Synechocystis* cells exposed to different stress factors (*Figure 1*), we were expecting to see differences in fluorescence intensity of the samples, which (in an ideal case) would be proportional to the concentration and fluorescence quantum yield of the pigments^50^. In order to separate the effects of IoR from concentration and functional state of PBs, we used time-resolved fluorescence spectroscopy with picosecond time resolution (*Figure 2AB*). Surprisingly, the reduction of PBs absorption was accompanied by an increase of the PBs fluorescence intensity, which is due to the increase of PBs fluorescence lifetime and, hence, quantum yield (*Figure 2C*). Since under normal conditions PBs fluorescence is quenched due to highly efficient excitation energy transfer (EET) to the chlorophylls of the photosystems^6,17^, we assume that the observed increase in PBs fluorescence lifetime is due to uncoupling of PBs from the photosystems. The decrease of the EET efficiency explains the characteristic changes in steady-state fluorescence spectra, in which the intensity of the PBs emission band (660 nm) significantly increases, while Chl emission at 685 nm decreases (*Figure 2D*) after exposure to IoR or during the space flight. Decrease of Chl fluorescence intensity is probably caused by a combination of several factors (i) decrease of the EET efficiency form PBs, (ii) nonphotochemical quenching and (iii) decrease of Chl concentration in photosystems. Uncoupling of PBs from photosystems also explains the increase of the initial level of fluorescence intensity (*F0*) observed in the fluorescence induction curves of the sample exposed to IoR (*Figure 2E*). We found that so-called photosynthetic efficiency of PS2 (*Fv/Fm*) significantly decreases with increasing radiation dose (*Figure 2F*), which could be partially explained by an increase of *F0*; however, activity of PS2 is definitely affected, since no J-P stages could be found at 120 kGy. Thus, under IoR, PS2 gets into Q_B_ non-reducing state. Since the value of *Fv/Fm* of cells which returned from space was close to zero, the space flight also inactivated PS2. In contrast, the activity of PS2 of the control sample, which stayed in the dark during the time of the space flight, remained high (*Fv/Fm =* 0.46), although the concentration of PBs antennas was significantly reduced (see *Figure 1C*).

**Figure 2.**
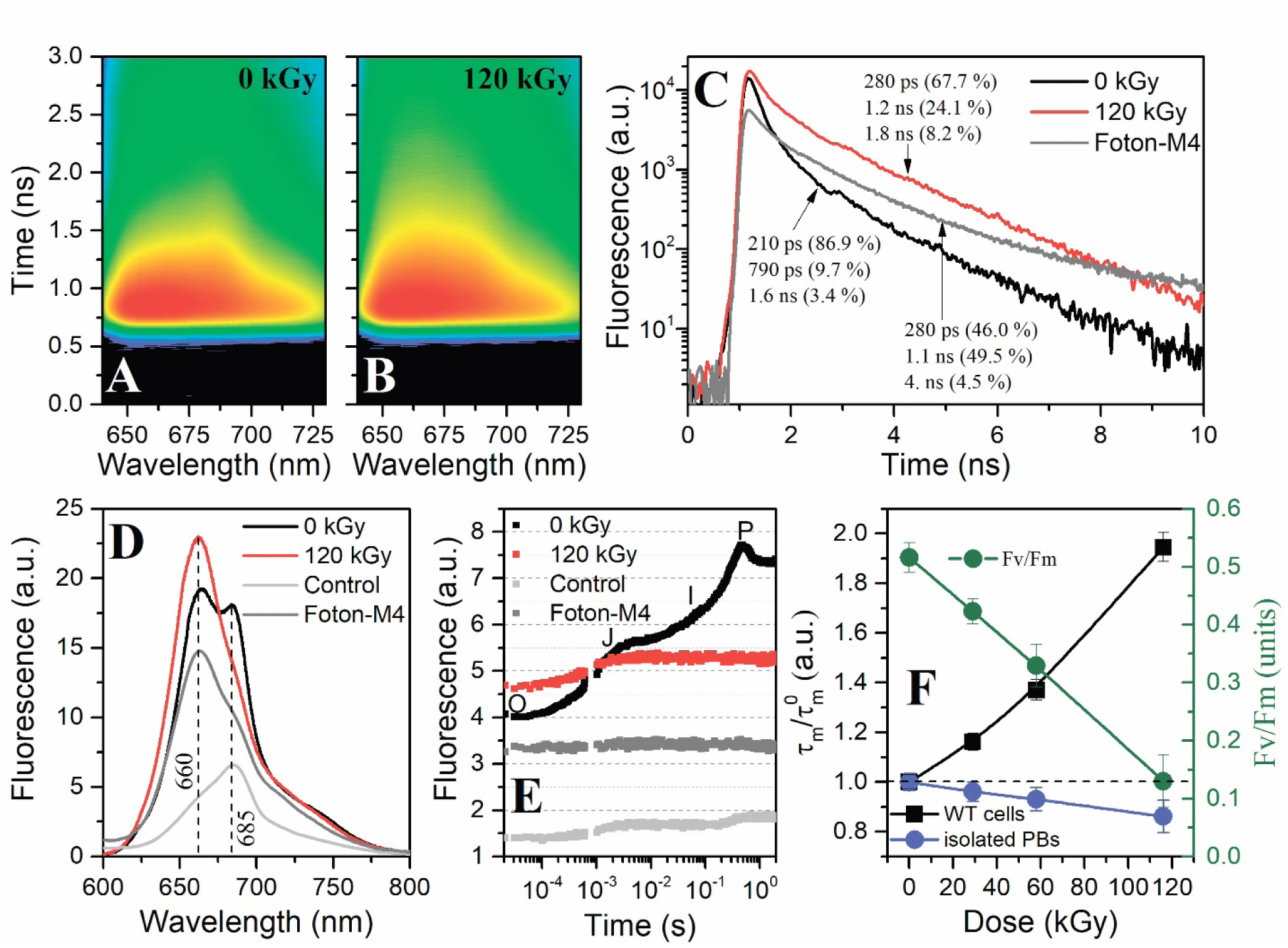
(**A**) and (**B**) – time-resolved emission spectra of *Synechocystis* sp. PCC6803 samples before and after exposure to 120 kGy of α-particles, respectively. The color represents the logarithm of the photon number in each time/spectral channel. Experiments were conducted at 25 °C. (**C**) – fluorescence decay kinetics of *Synechocystis* sp. PCC6803 samples before (black) and after exposure to 120 kGy of α-particles (red), and after Foton-M4 space mission (gray, scaled) at 660 nm emission. (**D**) – steady-state fluorescence spectra of *Synechocystis* sp. PCC6803 samples before and after exposure to 120 kGy of α-particles (black and red, respectively), Foton-M4 space mission (gray, scaled) and the control sample incubated in the dark for the duration of the space flight (light gray, scaled). (**E**) – fluorescence induction curves of *Synechocystis* sp. PCC6803 cells, color codes as in the previous panel. (**F**) – dose dependency of fluorescence lifetimes of *Synechocystis* sp. PCC6803 cells and isolated PBs, normalized to the corresponding values at 0 kGy. The right Y-scale shows the dose dependency of the Fv/Fm ratio calculated from fluorescence induction curves (see panel E).

The most surprising effect was observed in experiments with PBs isolated from *Synechocystis* cells. Since EET in PBs proceeds from blue (PC, emitting at 647 nm) to red pigments (allophycocyanines (APC) and terminal emitters, at ∼ 660 and 685 nm, respectively), we hypothesized that IoR may cause decoupling of the PBs rods (mainly phycocyanines) and cores (mainly allophycocyanines), which would result in an increase of the PC fluorescence quantum yield. If it was the case, the decomposition of PBs into individual parts would explain the observed increase of PBs lifetimes *in vivo*. In contrast, we observed a decrease of PC and APC lifetimes in isolated PBs under IoR (*Figure 2F*). This fact, together with the absence of a (significant) blue shift of PBs fluorescence emission *in vivo* (*Figure 2D*) indicates that, under IoR, PBs uncouple from the photosystems. However, PBs do not dissociate, since EET from PC to APC remains effective. This shows that IoR-induced uncoupling of PBs occurs in a controlled (and probably reversible) manner, similar to state transitions. Since it is highly unlikely for cyanobacterial cells to have a specific receptor for α-particles, we assume that PBs uncoupling from photosystems could be a general protection mechanism which could be triggered by multiple stress factors, and most probably ROS.

### 3.3 Ionizing radiation and ROS can provoke photoprotective mechanisms in vitro

It is known that the shape of the absorption spectrum, fluorescence lifetime and quantum yield of phycobilins is determined by interactions with the protein matrix, and denaturation of the protein causes significant reduction of the fluorescence lifetime^51,52^. If this happens when the phycobiliprotein is part of the PBs antenna, such a chromophore becomes a trap for excitation energy, which will be dissipated as heat. Since thermal dissipation of excitation energy is a major photoprotection strategy for all cyanobacteria, which are endowed with an OCP-dependent quenching mechanism of PBs fluorescence^31,53-55^, we decided to test how IoR affects the properties of phycobiliproteins and OCP individually.

The absorption characteristic for PC gradually decreases under exposure to IoR (*Figure S1A*). At 120 kGy, less than 5% of the initial absorption of PC remains in the region from 500 to 700 nm. Surprisingly, we found almost no changes in fluorescence lifetime (data not shown), although fluorescence intensity decreased tenfold, which indicates that the concentration of chromophores decreases. This is in a line with the observed gradual decrease of the intact protein bands at 17 and 22 kDa^12^ resolved by SDS-PAGE (*Figure S1B*). However, we expected to see new bands on the gel, representing shorter fragments of peptide chains, or a gradual decrease of the molecular weight of PC polypeptides, but this was not the case. Thus, IoR reduces the concentration of PC in solution, but there was no evidence for accumulation of phycobilins released into solution or of denatured protein with phycobilins still bound and solvent-exposed. The bleaching of PC pigments upon exposure to IoR could be explained by two major reasons: (i) direct impact of the α-particle and (ii) generation of ROS in traces of α-particles. Indeed, we observed gradual bleaching of PC in experiments in which ROS was generated by a photosensitizer (data not shown). However, the most interesting effect was found in experiments with OCP.

**Figure S1.**
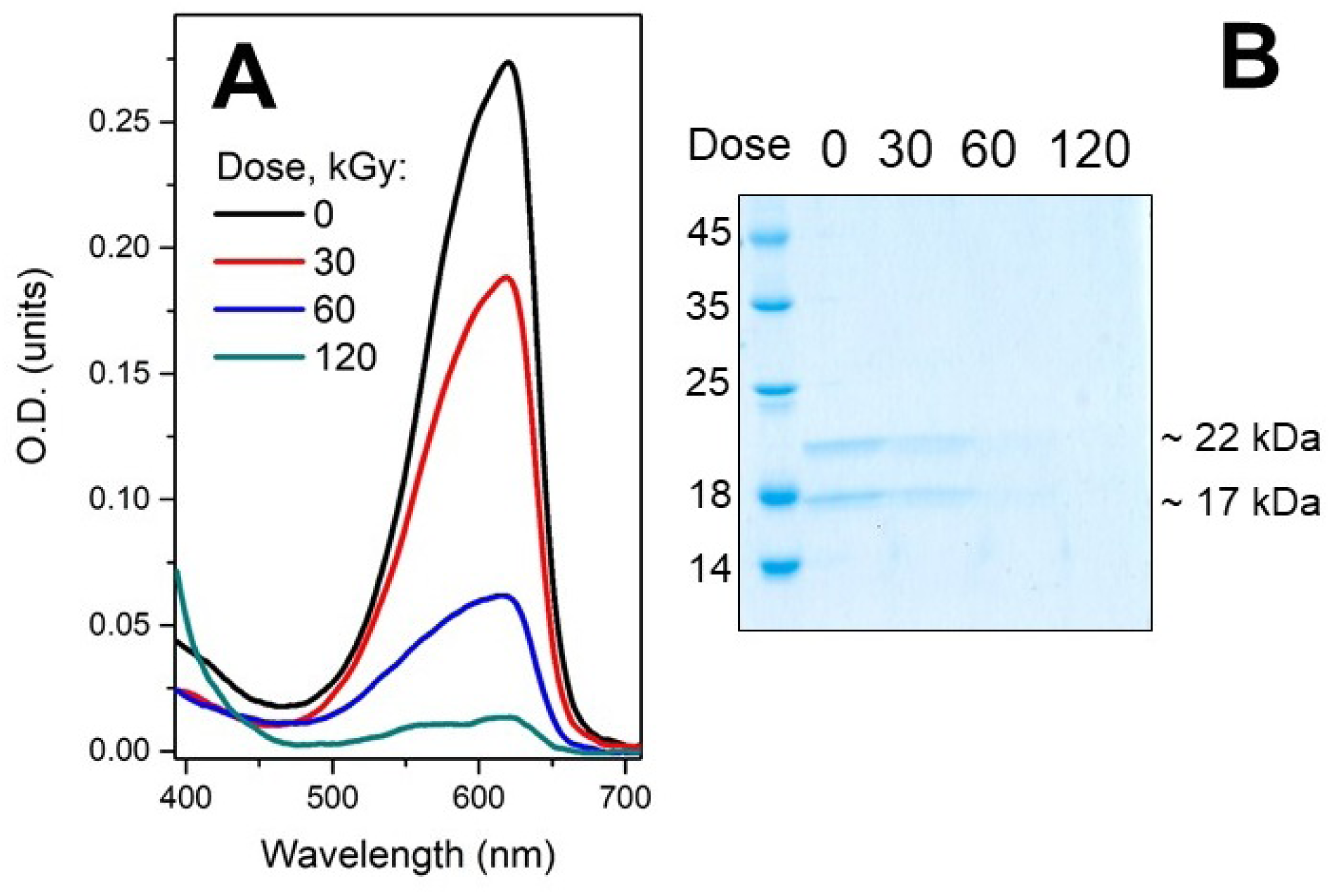
(**A**) absorption spectra and (**B**) SDS-PAGE of PC of PC after the exposure of the sample to different doses of α-particles. Two major bands could tentatively be assigned to individual α and β polypeptides of PC.

OCP is bleached upon generation of ROS *in vitro*, but during this process, the protein converts into a state with the red-shifted absorption spectrum (*Figure 3*). Upon illumination of OCP by the same red-light source in the absence of AlPCs as photosensitizer, we observed no changes in the absorption spectrum (data not shown). However, in the presence of AlPCs, which are characterized by a high-yield ROS production^56^, all absorption bands of OCP in the visible region of the spectrum gradually disappear (*Figure 3A*). The changes in O.D. below 500 nm (*Figure 2C*) could well be approximated by a single exponential decay component. At the red flange of OCP absorption the time course is more complex and even exhibits a rising part at the beginning of the ROS experiment. An O.D. increase in this spectral region (500 – 600 nm) is characteristic for accumulation of the red active signaling state of OCP^45,54^. According to our results, ROS-induced increase of O.D in 570 region is equivalent to 20 % of changes upon the photoactivation of OCP (*Figure 3B*). Thus, ROS stress induces the transition of OCP from its stable orange state into the active red form which is able to bind PBs and induce fluorescence quenching. It is known that OCP could be activated “chemically” by high concentrations of chaotropes, for example 1.5 M NaSCN^42,57^, thus destabilization of the native OCP structure is expected to trigger conversion into the red state, equivalent to photoactivation. Along these lines, nonspecific action of ROS can also somehow destabilize and thereby activate OCP. This observation is important for understanding of photoprotection in cyanobacteria, since it shows that protective mechanisms could be activated not only by strong light, but a variety of stresses in general.

**Figure 3.**
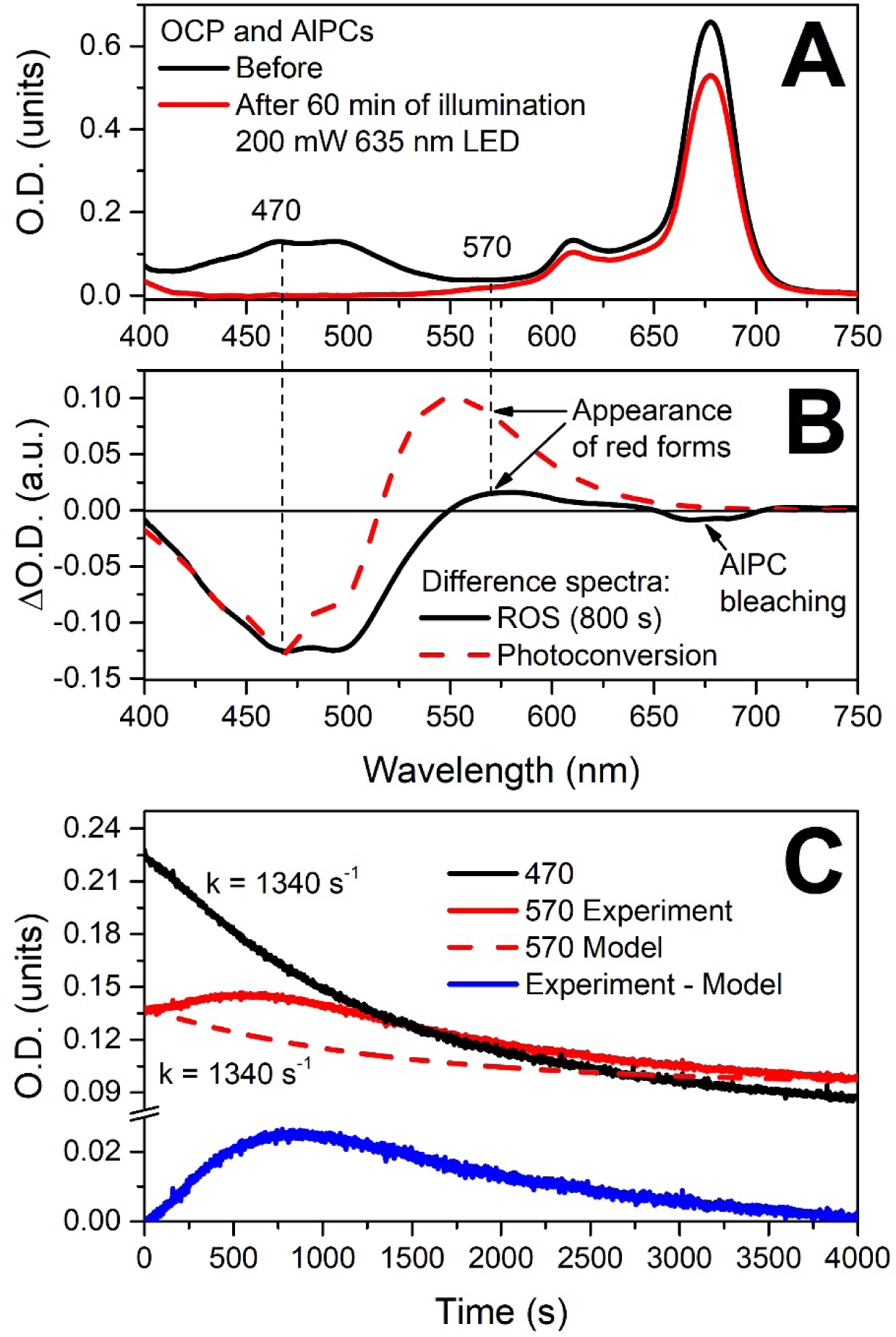
Bleaching of the carotenoid chromophore of OCP under photoinduced ROS production by aluminum phthalocyanine (AlPC). (**A**) – absorption spectrum of an OCP/AlPC mixture before and after illumination of the sample by red light triggering ROS production by AlPC. (**B**) – difference spectrum indicating accumulation of red forms of OCP under ROS production (black line) in comparison to photoactivation of OCP by actinic blue light (red dashed line). (**C**) – kinetics of OCP bleaching, measured as changes of O.D. at 470 and 570 nm. The blue curve represents kinetics of the red OCP accumulation (see text for details).

## 4. Discussion and Conclusions

Both, the examination of panspermia hypothesis and the exploration of possibilities for mankind to leave the Earth and to send seeds of Life to other planets requires estimation of the influence of space flight conditions on major physiological functions and the viability of different species in general. Since relatively primitive life forms were able to successfully survive in harsh conditions during the early stages of Earth’s history (and even today, see e.g. *Deinococcus radiodurans^20^*) and are considered as possible pioneers in colonization of other planets, their safe transportation is a challenge for future space missions. Among the stress factors, cosmic rays represent a major concern due to their ionizing ability. Thus, the evolution of the species before formation of the protective shields of the ozone layer and ionosphere required the development of radioprotective mechanisms.

In this work, we analyzed the effects of space flight during the Foton-M4 mission and of direct irradiation by α-particles with energies of about 30 MeV on the functional organization of the photosynthetic apparatus and primary stages of photosynthesis of the cyanobacterial strain *Synechocystis* sp. PCC6833. In order to study the action mechanism of irradiation with α-particles, we also performed experiments on the components of the photosynthetic apparatus on different levels of organization from whole cells over supercomplexes of light-harvesting proteins down to individual protein components. We found that the light-harvesting antennas - phycobilisomes - are extremely sensitive to IoR *in vitro*. The differences in effects observed *in vivo* and *in vitro* indicate that cyanobacteria still have general protection mechanisms and durability which probably allowed them to survive and evolve during early stages of the Earth’s history, long before the Great Oxygenation Event. We found that *Synechocystis* sp. PCC6833 can uncouple PBs from photosynthetic membranes (*Figure 2*), which could be a first step to prevent radiodamage and overexcitation of photosystems. This event requires mobility of the PBs antennas, which is also observed during state transitions; however, in this case, mobilization is rather triggered by ROS stress than by the cellular redox state. We also found evidence that ROS may activate OCP-dependent photoprotection. Such a combination of protective mechanisms allows to prevent damage of photosystems and to keep PBs quenched while being detached, in order to prevent additional ROS production. Probably, water-soluble carotenoid-containing proteins from the OCP superfamily could be involved in such regulatory processes. Some of the various types of helical carotenoid proteins (HCPs), which are homologs of the OCP N-terminal domain, do not interact with PBs and have no function assigned yet. It is conceivable that they could be remnants of ancient protective mechanisms, or that the site of their interaction with PBs is not available in the native state of the PBs. These hypotheses require more experimental scrutiny, since at the moment, even for the most comprehensively studied OCP from *Synechocystis*, the exact site of interaction with PBs is unknown ^*58*,*59*^.

The production of huge amounts of proteins to sacrifice them in order to salvage DNA from damage is thought to be a general strategy for microorganisms to survive at high levels of IoR. Species like *Deinococcus radiodurans* are living proofs that this simple strategy works perfectly^60^. We can assume that today’s cyanobacteria like *Synechocystis* inherited variant relics of this protective mechanism even in the absence of excessive IoR stress due to the involvement of PBs in photosynthesis, which gave them numerous other competitive advantages. It is not clear whether PBs had both radio- and, more general, ROS-protective as well as light-harvesting functions from the beginning, since we know little about the evolution of PBs. However, we can postulate that in the absence of IoR as a selection factor both of these roles became advantageous, since both, optimization of light-harvesting and protection against ROS stress, are persistent issues for successful propagation of a species. Perhaps some of these questions could be answered by considering the reasons for the loss of PBs in algae and higher plants from the perspective of functional diversity of these megacomplexes.

After the space mission and exposure to high doses of IoR, *Synechocystis* sp. PCC6833 cells were able to grow autotrophically again and produced cultures with spectral and functional characteristics indistinguishable from the control samples. As approved space veterans, they are now introduced into the collection of microorganisms of Biological Faculty of Lomonosov Moscow State University.

## Acknowledgements

This work was supported by the Russian Foundation for Basic Research (projects 16-34-00394, 18-04-00554). This work was supported by Russian Ministry of Education and Science (MK-951.2018.4). T.F. and acknowledge support by the German Federal Ministry of Education and Research (WTZ-RUS grant 01DJ15007) and the German Research Foundation (Cluster of Excellence “Unifying Concepts in Catalysis”).

